# Deep and Quantitative Proteomic Profiling of Low Volume Mouse Serum Across the Lifespan

**DOI:** 10.1101/2025.06.24.661372

**Authors:** Amit K Dey, Bradley Olinger, Mozhgan Boroumand, Maria Emilia Fernandez, SLAM investigators, Simonetta Camandola, Nathan L Price, Rafael de Cabo, Nathan Basisty

## Abstract

Assessing and validating circulating biomarkers is essential for the development of pre-clinical biomarkers that predict biological aging and aging-phenotypes in mice. However, comprehensive proteomics of serum, especially in longitudinal mouse studies, are limited by low volumes of samples. In this study, we develop a workflow for comprehensive and quantitative proteomic analysis of low volume mouse serum and demonstrate its utility and performance in identifying and evaluating key associations with aging phenotypes. Notably, a nanoparticle (NP)-based serum processing workflow coupled to mass spectrometry (MS) increases proteomic coverage by 3 to 6-fold across a range of volumes and provides a quantitative and reproducible (CV < 10%) pipeline for NP-based studies. In a study of 30 mice (aged 12, 24, and 30 months), we uncovered 3992 protein groups across all samples (2235 on average) in 20 µL of serum and highlight novel insights into aging-associated changes in serum and associations with glucose and body composition. With 1 µL additional serum, a 48-cytokine assay quantified 39 additional proteins not identified by MS. This study establishes a powerful workflow that enables deep quantitative proteomics of biologically relevant proteins in volumes feasibly obtained from mice (21 µL of serum) and presents fundamental insights into the aging serum proteome.

## INTRODUCTION

Blood-based specimens such as serum are rich in proteins originating from diverse organs that offer valuable clinical potential [1, 2] and ease of repeated collections with minimal invasiveness. Work from our group and others have highlighted the blood proteome as a source of biomarkers associated with aging-related phenotypes, including changes in body composition and glucose, in human clinical studies [3-7]. The full potential of using the circulating proteome to measure aging and predict health in humans has not been fully realized due to the practical challenges of longitudinal sampling across the human lifespan [8]. However, mouse models, due to their short lifespans, are useful surrogates for rigorously exploring longitudinal changes in the circulating proteome throughout lifespan and formulating strategies for quantifying age-related organismal declines with protein biomarkers in humans [9-11]. Comprehensive serum biomarker assessment is lacking in mice for aging studies due to 1) the dynamic range of protein concentrations in blood and 2) limited volume compared to human blood specimens [12]. Mass spectrometry-based proteomic analysis of serum is technically challenging due to a dynamic range of protein concentrations in blood spanning over 10 orders of magnitude [13], presenting a formidable obstacle in detecting and quantifying proteins comprehensively, particularly low abundance proteins that tend to be promising biomarker candidates. Secondly, deep proteomic studies are not feasible in mouse studies with limited blood volume, particularly when blood is collected longitudinally or is needed for multiple assays. A truly rigorous assessment of aging biomarkers requires a comprehensive and longitudinal assessment of aging-related phenotypes in parallel with sample collections to make associations of protein changes with phenotypes and phenotype trajectories.

Over the past decade, LC-MS methodologies, alongside advancements in sample preparation strategies and computational tools, have significantly improved depth, sensitivity, and throughput for biomarker discovery [14]. Nanoparticle (NP)-based protein corona workflows have recently emerged as a promising method to reduce sample dynamic range and improve the detection of low-abundance proteins [15]. In doing so, NP-based enrichment coupled with LC-MS like data-independent acquisition (DIA) permits comprehensive, reproducible, and deep plasma proteome coverage in blood-based samples, facilitating unbiased biomarker discovery and insights into biological pathways at the scale required for biomarker studies [16-18], enabling the reproducible identification and quantification of upwards of 6,000 proteins in human plasma. Despite its impressive depth, MS struggles with detection of cytokines and other key players in “inflammaging” [19], owing to several factors: a limited number of unique tryptic peptides present in small proteins such as cytokines, the low endogenous concentrations of cytokines (in the pg/mL range), transient secretion, and short half-life [20]. Sensitivity is further compromised by matrix interferences and inefficient peptide ionization [21]. To overcome limitations in cytokine detection by MS, complementary affinity-based methods such as proximity extension assays (PEA) offer multiplexed quantification of circulating cytokines, detecting up to 48 cytokines with reasonable volume requirements, 1 µl of mouse serum. Multiple studies highlight the potential benefits of integrating untargeted LC-MS with targeted immunoassay PEA or aptamer-based assays offering valuable complementarity for a biologically relevant gain in proteome coverage [22-25].

The present study aims to demonstrate the feasibility of performing comprehensive proteomics on small-volume collections in mice compatible with longitudinal studies within the framework of NIA’s Study of Longitudinal Aging in Mice (SLAM) [11], an effort to collect repeated measures on functional, phenotypic, and biological variables across lifespan to rigorously characterize normative aging in mice. We utilize a subset of SLAM samples to demonstrate the feasibility and capabilities of an automated NP-based sample processing workflow to perform comprehensive and reproducible aging biomarker studies in limited volumes of serum on an automated platform that can be applied at the scale of SLAM. We develop a quantitative pipeline and benchmark the performance of NP-based MS for accurate, quantitative, and reproducible biomarker studies in low-volume mouse serum. We also combine untargeted LC-MS with targeted PEA assays, highlighting the complementarity of a combined approach in identifying and quantifying cytokines while demonstrating the strengths and limitations of each approach. We evaluate and optimize the data analysis pipeline required for accurate quantitation of proteins from nanoparticle fractions, benchmarked to ground truth samples. Finally, we apply this approach to a study of aging and aging-related clinical outcomes in adult, old, and geriatric mice to gain unprecedented depth and novel insights into aging through the circulating proteome, demonstrating the feasibility of conducting deep and quantitative serum proteomics and biomarker discovery in sample-limited mouse studies.

## RESULTS

### Qualitative and quantitative evaluation of a nanoparticle-coupled serum proteomics workflow

A multi-component study design was applied to develop a comprehensive proteomic quantitation method compatible with low-volume serum, and to validate its accuracy and ability to identify circulating biomarkers (**Fig 1A**). A combination of nanoparticle-coupled mass spectrometry workflow (**Fig 1B**) and affinity proteomic assays (**Fig 1C**) were leveraged to detect and quantify serum proteins for an assessment of aging and pre-clinical biomarkers (**Fig 1D**) in a mouse cohort. To initially evaluate the feasibility of performing comprehensive proteomics in sample-limited mouse serum, we adapted an automated nanoparticle-based sample processing workflow in a range of serum volumes. A feasibility study comparing the number of proteins and peptides identified in 10, 20, 50, and 100 µL of pooled mouse serum showed 2 to 4-fold improvement in protein depth compared with non-volume limited neat digests (**Supplementary Fig. 1A**). This finding aligns with previous observations indicating that nanoparticles can reduce the dynamic range of protein concentrations in serum and improve protein depth [16]. To qualitatively assess the effect of serum preparation methods, we compared the proteins identified in serum collected by two common methods: 1) Clotting for 30 min at room temperature and 2) collection in Sarstedt Z-gel tubes. We identified 1700 and 1681 protein groups in each respective method, of which over 90% were shared (**Supplementary Fig. 1B**), suggesting the method of serum preparation does not substantially impact the performance of the method. For all downstream analysis, method 1 was used for serum collection.

**Figure 1:**
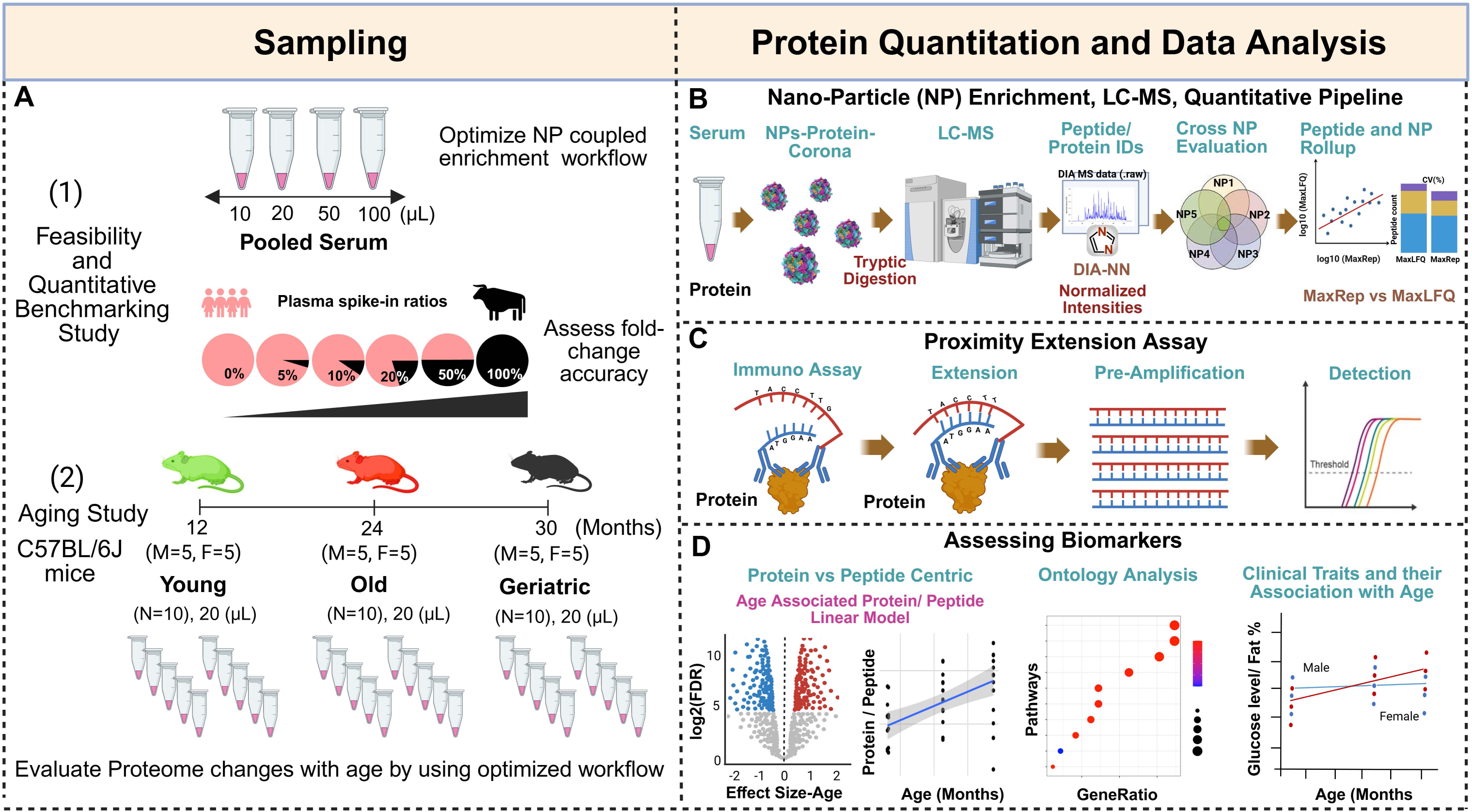
Overview of study design for a comprehensive and quantitative proteomic study in mouse serum. A) Schematic summary of study design with two phases: 1) a feasibility and quantitative benchmarking study and 2) an aging study. In phase 1, to initially assess the feasibility of this workflow at low volumes, optimized nanoparticle enrichment and LC-MS workflow with pooled mouse serum was conducted with four different volumes (10, 20, 50, and 100 µL), and compared to traditional neat serum digest (no enrichment) processed in parallel. To assess quantitative accuracy, a benchmarking study was conducted on defined ratios serum from two organisms (bovine and human). In phase 2, to demonstrate the power of the workflow in a biological study, an aging study of 30 mice was conducted at three ages: 12 (young), 24 (old), and 30 (geriatric) months. B) The automated serum-nanoparticle processing and digestion were performed using a Proteograph (Seer, inc) workflow. Peptide data was acquired by nLC-MS/MS and analyzed with DIA-NN in a library-free mode. NP-peptide enrichment was assessed by comparing two generic label-free quantification algorithms, MaxRep and MaxLFQ. C) A panel of 48 mouse cytokines were measured using proximity extension assay (O-link, Thermo Scientific). D) Biomarkers and pathways associated with aging and clinically relevant phenotypes were assessed.

To assess whether this workflow enhanced the detection of biologically relevant proteins, we assessed proteomic coverage of known aging-associated biomarkers and processes. Cellular senescence is a widely accepted hallmark of aging, and senescence associated proteins in circulation are clinically relevant biomarkers in aging studies [3, 4, 7]. In the feasibility study, we detected multiple canonical senescence-associated secretory proteins spanning the dynamic range of the sample, including Igfbp’s 2/3/4/5, Timp’s 2/3, Thbs1, and Mmp’s 2/8, among other known markers (**Supplementary Fig. 1C**). For a more comprehensive assessment in the aging study, we evaluated the detection of a recently described *in vivo* mouse senescence signature, SenSig, in serum [26]. Compared with neat serum, where 45 SenSig proteins were identified, 136 and 209 SenSig proteins were identified in NP processed samples or across all samples in the aging study (**Fig. 2H**), respectively. These results demonstrate that this workflow substantially increases the detection of biologically relevant, low abundance proteins.

**Figure 2:**
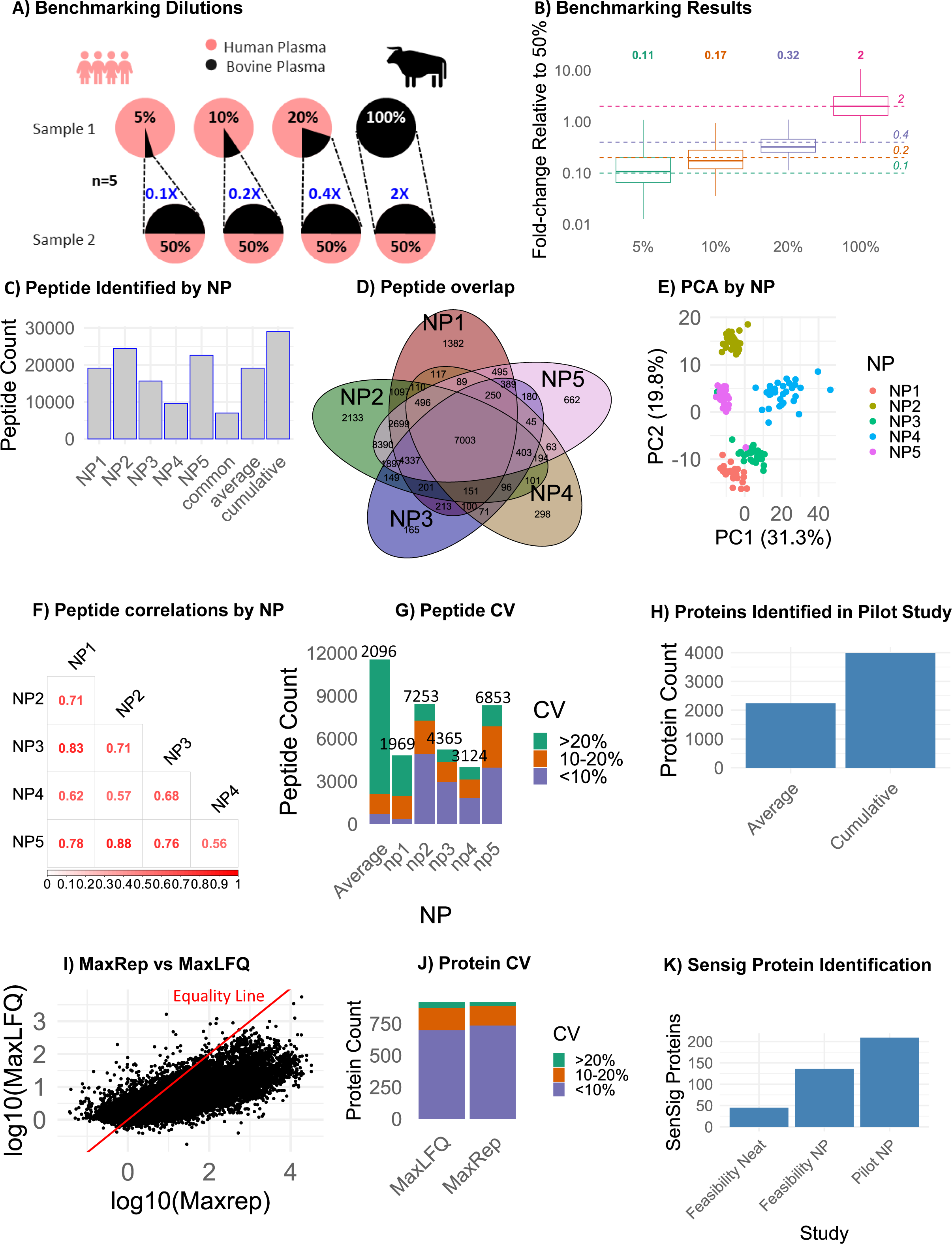
Qualitative and quantitative evaluation of proteomic data generated NP-coupled MS workflow. A) A quantitative benchmarking study was performed using five fixed ratios of human and bovine serum (n=5 each). B) Peptide abundance ratios in each dilution versus the 50% bovine serum sample. Dotted lines depict the expected ratios. C) Peptides identified in at least one sample by NP. D) Proteins quantified in at least one sample are shown for each NP. E) Principal Component Analysis was used to show clustering patterns between nanoparticles on the peptide level. F) Spearman correlation values for peptide intensities between NP. G) Coefficient of variation (SD/Mean abundances) is shown by NP on the peptide level for 3 triplicate samples. For each NP, only those peptides detected in all 3 samples were used for calculation. For combined, intensities were added between all NP and proteins detected in all 3 samples (regardless of which NP) were kept for analysis. H) Protein identifications in mouse aging study. I) Correlation of protein intensities calculated by the MaxRep versus MaxLFQ approaches. J) Protein CVs in the MaxRep and MaxLFQ quantitation pipelines. K) Number of SenSig proteins detected in neat serum and serum processed by the NP workflow.

A measurement is quantitative when the relative change in the measured signal reflects the change of the peptide quantity in the original samples. Given the known effect of the nanoparticle-coupled MS workflow on the dynamic range of proteins in a sample, it is critical to determine whether the samples produced by this workflow preserve the quantitative differences in the original samples. To evaluate the ability of NP processing workflow to accurately quantify peptide abundance, we conducted a benchmarking study similar to previously described matrix-matched calibration curves and LFQ-Bench studies [27, 28]. Briefly, we produced benchmarking samples containing known ratios of human and bovine serum (**Fig. 2A**), including 19:1 (5% bovine), 9:1 (10% bovine), 4:1 (20% bovine), 1:1 (50% bovine), and 0:1 (100% bovine) ratios (n=5 each). Each sample/dilution was processed through the standard NP workflow. In this way the accuracy of the NP workflow can be assessed by comparing reported peptide abundances between samples with known ratios. The fold change in peptide intensities were calculated for all dilutions relative to the 50% bovine samples. The measured peptide ratios were similar to the expected ratios (**Fig. 2B**), with the median of the 5% and 100% bovine samples being within 0.01 of the expected ratios and the expected ratios of the 10% and 20% bovine sample groups within their respective interquartile range. These findings suggest that the NP-workflow accurately depicts the relative changes of the original serum samples.

### Evaluation and optimization of NP-coupled MS in low-volume mouse serum study of aging

To evaluate the performance of the nanoparticle-coupled MS workflows in identifying serum biomarkers, we applied this method in a subset of 30 C57BL/6J mice from SLAM. We examined 20 µL serum samples per mouse at three ages: young (12 months, n=10), old (24 moths, n=10) and geriatric (30 months, n=10), with each group composed of 50% females (**Fig. 1A**, “Aging Study”). Following nanoparticle processing, captured proteins were digested using trypsin/ Lys-C and the resulting peptides were analyzed by LC-MS in data-independent acquisition mode (DIA). DIA data were analyzed using DIA-NN database search algorithm within the Proteograph Analysis Suite (PAS).

Across all 30 mice, approximately 10,000-25,000 peptides were detected within each NP fraction, and 28,976 unique peptides were cumulatively detected (**Fig. 2C**). In rank order of the number of peptides identified, 24457 peptides were detected using NP2, 22592 peptides with NP5, 19129 peptides with NP1, 15650 peptides with NP3, and 9587 peptides with NP4. A total of 7003 peptides were commonly detected across all nanoparticles, and 1382, 2133, 165, 298, and 662 peptides were uniquely identified, respectively, in NPs 1-5 (**Fig. 2D)**. Principal component analysis revealed clustering of NP by peptide expression levels **(Fig. 2E)**. Among peptides quantified in all NPs, moderate to high correlations in peptide expression levels by NP were observed, such as between NP1 and NP3 (rho = 0.83) and NP2 and NP5 (rho = 0.88), (**Fig. 2F**). Total peptide counts and coefficient of variation (CV) from technical triplicates with 20 µL serum volume showed that NP2 and NP5 exhibited the highest peptide detections, corroborating the NP ranks observed in Fig. 2C (**Fig. 2G**). However, NP2 demonstrated the greatest precision in peptide identification with a median CV of 9% (**Supplementary Fig. 2A)**, with over half of the total identified peptides having a CV of less than 10%. As expected, the CVs of the cross-NP average are the highest, consistent with differences in the relative affinity of each nanoparticle for any given peptide. Overall, this contrast between nanoparticles is consistent with previous studies [15, 29] suggesting that distinct nanoparticles provide heterogeneous and reproducible surface chemistries that facilitate the formation of distinct corona compositions.

### Protein-level quantification from NP fractions: Comparing roll-up approaches

To develop an optimal pipeline for quantitation of proteins with independent peptide measurements across distinct nanoparticle fractions, we compared two established rollup approaches: MaxRep and MaxLFQ. For MaxRep, for any given peptide, the greatest observed intensity across the nanoparticles is used for quantitation. Subsequently, MaxLFQ, a previously described peptide rollup approach [30], was applied to conduct peptide rollup. In the MaxLFQ-only approach, the rollup is conducted across both the nanoparticle and peptide level for all observations, as previously reported on this platform [29]. Using either method, a total of 3992 proteins were quantified cumulatively (2235 protein groups on average) (**Fig. 2H**), with a median of 4 peptides mapped to each protein (**Supplementary Fig. 2B**) and a median of 63 distinct peptide-NP measurements were observed per protein (**Supplementary Fig. 2C**). MaxRep and MaxLFQ methods were significantly correlated (correlation = 0.77, p-value: <0.00001), although the correlation is weakest among high intensity observations (**Fig. 2I**). To assess precision of each protein quantitation workflow, we analyzed proteins quantified by five NPs across technical triplicates of 20 µL serum samples. The median protein-level CVs were 0.0567 and 0.0613 for MaxRep and MaxLFQ, respectively, and the proportion of proteins below 10% CV was modestly higher in MaxRep (**Fig. 2J**). Overall, these findings suggest that the NP-based workflow, particularly the MaxRep pipeline, produces quantitative and reproducible protein measurements. A consistently higher number of quantified proteins observed in NP-processed samples compared with the standard (neat) samples, irrespective of data processing algorithms or nanoparticle fraction, is consistent with a reduction in the protein dynamic range concentration in protein coronas generated by NP-processing.

### Identification of cytokines commonly missed by MS

While a large number of proteins, including inflammation and senescence-associated proteins (**Fig. 2K, Supplementary Fig. 1C**), were detected following the NP workflow, prototypical inflammatory cytokines, such as IL-6, were notably not detected, consistent with previous MS studies [4, 31]. Given the important role of cytokines in a range of diseases and aging, including as a key bioactive component of the senescence-associated secretory phenotype, we sought to enhance the proteomic dataset with a complimentary targeted approach requiring minimal additional serum. Using a targeted 48-cytokine panel, 43 inflammatory cytokines were quantified in 1 µL of additional serum from each mouse (**Fig. 3A, Supplementary Table 1**). While heterogeneity can be seen between biological replicates at any age, a general increase in abundance of proteins across the panel with age is apparent (**Fig 3A**), and age and sex-dependent separation is apparent by principal component analysis (**Fig 3B**). Notably, only four proteins were detected by both MS and the cytokine panel (**Fig. 3C**), demonstrating the complementarity of this approach to detect biologically important cytokines that are missed by MS. To assess the agreement between the MS and antibody-based assays, we compared the data for IL16, which contained sufficient datapoints in both datasets for correlation. IL16 quantification was significantly correlated between the two approaches (p = 0.0085, correlation = 0.7455, **Fig. 3D**), confirming the MS-based quantitation of this protein.

**Figure 3:**
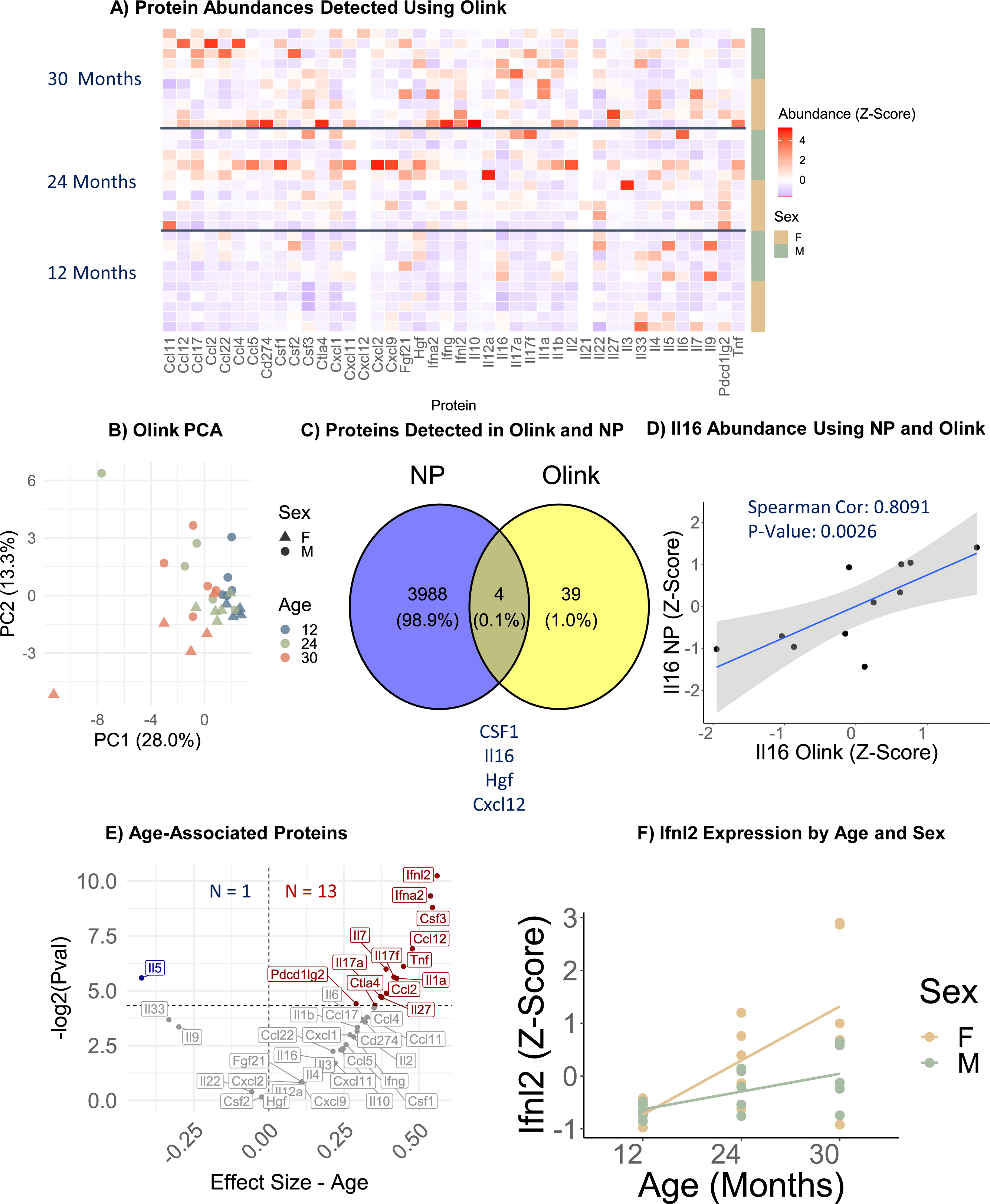
Age-related proteomic changes measured with proximity extension assays. A) Heatmap of scaled protein abundance for all proteins measured with PEA in the aging study (n=30 mice). B) Principal component analysis of protein abundance by age and sex. C) Proteins detected using both NP-MS and PEA methods. D) Correlation of IL16 levels measured by NP-MS and PEA. E) Pearson modeling revealing age-associated proteins measured by PEA using the formula: Protein ∼ Age + Sex. F) Ifnl2 levels by age in males and females.

To identify linearly age-dependent cytokine changes in circulation, linear modeling was performed of cytokine changes with age, adjusting for sex as a covariate. Notably, of 34 increasing cytokines, 13 cytokines significantly linearly increased, and one cytokine, interleukin-5 (IL5), significantly decreased with age (**Fig. 3E**), and interferon lambda-2 (Ifnl2) was the most significant change (**Fig. 3F**). Overall, these results highlight the complementarity of this approach to identify biologically relevant cytokine changes in 1 µL of serum.

### Age-associated changes in the serum proteome

We next examined circulating proteomic signatures of age from the NP-coupled MS workflow in a subset of samples from the SLAM. We selected proteins that had 2 or more mapped peptides (N = 2662 proteins) for subsequent analyses and ensured that each protein was present in at least three replicates from each age group and both sexes. All numerical protein data associated with the MS analysis is available in **Supplementary Table 2**. Principal component analysis revealed that samples were clustered by sex and modestly by age (**Fig. 4A**), whereas 73 proteins (54 increasing and 19 decreasing) showed statistically significant (FDR ≤ 0.05) linear association with age after adjustment for sex (**Fig. 4B**). The protein with the most statistically significant and positively age associated was Thioredoxin Domain-Containing Protein 5 (Txndc5), and Myosin light chain 3 (Myl3) . Other top significantly positively age-associated proteins include Actin Related Protein 2/3 Complex, Subunit 5 (Arpc5), Legumain (Lgmn), and Lectin Mannose Binding 1 (Lman1), among others (**Fig. 4C**). To identify non-linear proteomic changes across the lifespan, t-tests were performed to compare proteomic abundances between the young (12 months), old (24 months), and geriatric (30 months) mice. Notably, many proteins were differentially expressed at one life stage and not the other (**Fig. 4D**). Others were elevated at the 24-month stage and lowered at 30-months, while others showed the reverse trajectory (**Fig. 4E**). 189, 186, and 195 proteins in the old versus young, geriatric to old, and geriatric to young were upregulated, while 92, 165, and 148 proteins were downregulated in each, respectively (**Fig. 4F)**. Gene set enrichment analysis of the proteomic changes between the 12-month and 24-month groups revealed that gliogenesis, and immunity-related pathways are enriched (**Fig. 4G**), while the same analysis of the proteomic changes between the 24-month and 30-month mice identified metabolic processes to be enriched (**Fig. 4H**). Collectively, these results highlight the non-linear and stage-specific changes that occur in the circulating proteome with age.

**Figure 4:**
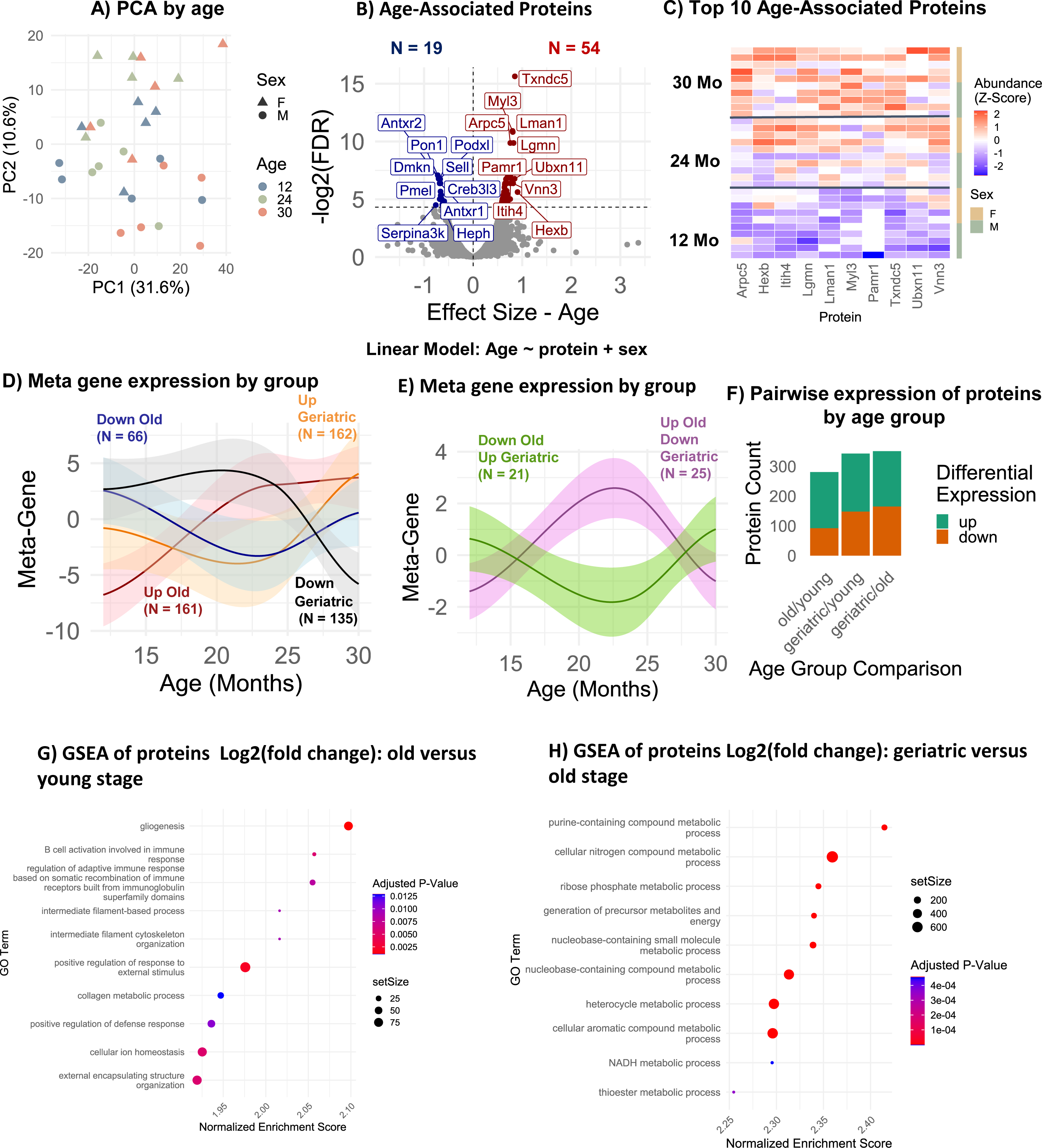
Age-associated changes in circulating proteins measured by NP-MS. A) Principal component analysis of quantified proteins by age and sex. B) Pearson linear modeling to identify age-associated proteins, using the model protein ∼ age + sex. Only proteins present in at least one sample from each age and sex group and with at least 2 peptides were examined (N = 2662). C) Scaled abundance of the top 10 linearly increasing age-associated proteins. D) Meta-gene expression of protein groups with stage-specific abundance changes with age. E) Meta-gene expression of protein groups with opposing stage-specific abundance changes. F) Differentially expressed proteins between young, old, and geriatric samples. Counts of proteins showing differential expression (p-values < 0.05) are shown at each stage. G) Gene Set Enrichment Analysis of protein abundance changes from ages 12 to 24 months (Biological Process). H) Gene Set Enrichment Analysis of protein abundance changes from ages 24 to 30 months (Biological Process).

### Sexual dimorphism of age-related changes in circulating proteins

Next, proteins that are differentially expressed by sex were identified using Pearson linear modeling in the 30-sample cohort using the NP-coupled MS workflow, which revealed 20 proteins to be upregulated in males, while 61 proteins were upregulated in females (**Fig. 5A**). Conducting the same analysis using the targeted cytokine panel, 5 proteins were significantly higher in males, including Ccl22, Il6, Hgf, Ccl17, and Cxcl1, while 2 were higher in females, including Pdcd1lg2 and Il7 (**Fig. 5B**). Using the proteins identified in the NP-coupled MS workflow, overrepresentation analysis of proteins elevated in males revealed that these proteins were implicated in the plasma membrane (**Fig. 5C**), while those elevated in females were implicated in peptidase activity, among other functions (**Fig. 5D**). Pearson linear modeling was similarly used to identify differences in proteomic trajectories with age that vary by sex by including an age-sex interaction term within the model, which identified 20 proteins that showed a stronger association with age in males compared with females, and conversely 52 proteins that showed the reverse association (**Supplementary Fig. 3A**). Interestingly, some proteins such as Col15a1 had a similar abundance between both sexes at early ages, but diverged significantly later in life (**Supplementary Fig. 3B**), whereas other proteins, such as Serpina1e, were differentially expressed by sex and maintained this contrast throughout the lifespan (**Supplementary Fig. 3C**). These findings suggest that this workflow is sufficiently to detect proteomic changes that emerge both by age and sex across the murine lifespan.

**Figure 5:**
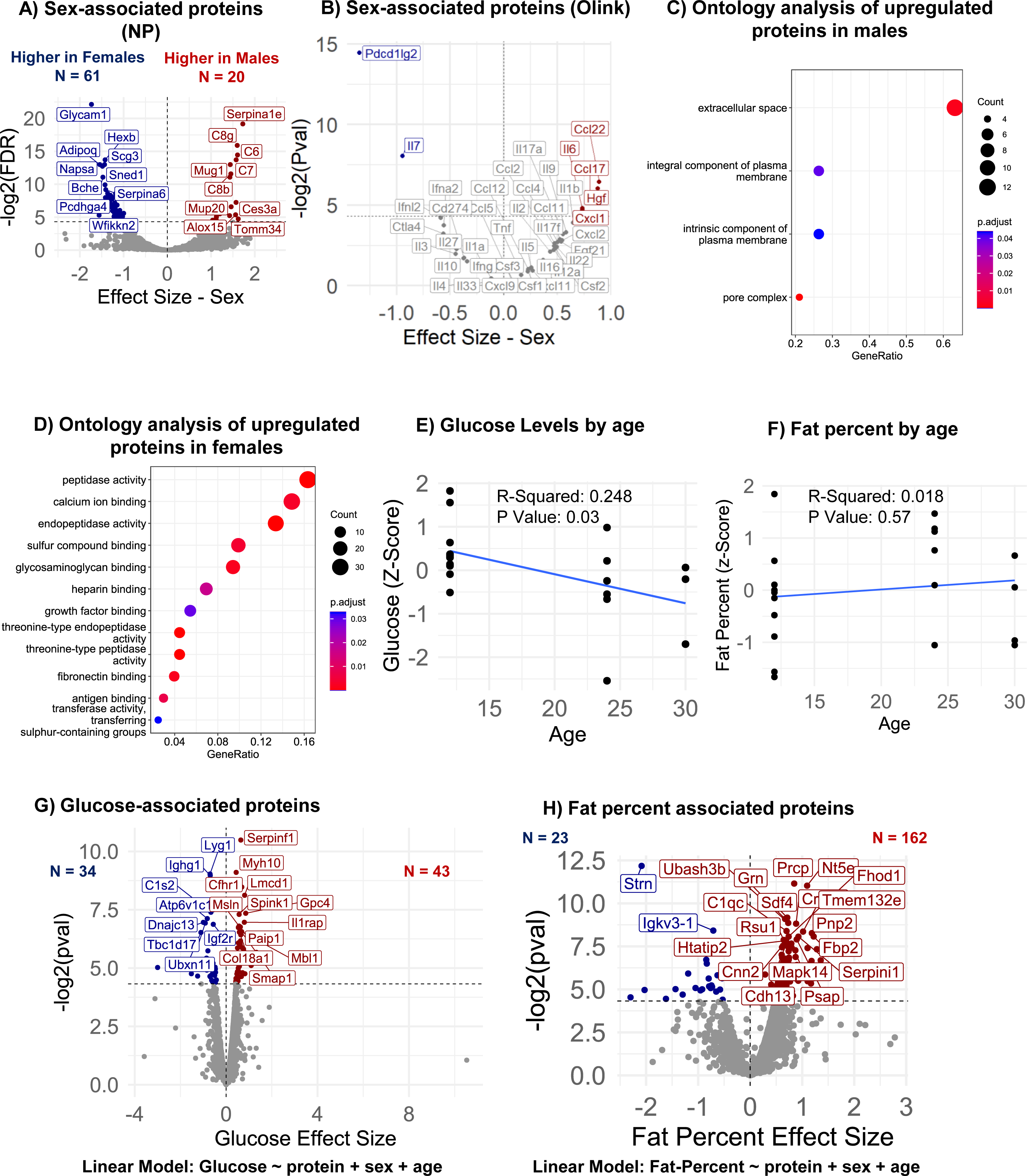
Circulating protein associated with sex and clinical traits. A) Pearson linear modeling to identify sex-associated proteins measured by the NP-MS workflow, using the model protein ∼ sex + age. B) Pearson linear modeling to identify sex-associated proteins measured by PEA workflow, using the model protein ∼ sex + age. C) Overrepresentation analysis of proteins upregulated in males (Cellular Component). D) Overrepresentation analysis of proteins upregulated in females (Molecular Function). E) Glucose levels with age. F) Total body fat percentage with age. G) Pearson linear modeling to identify glucose-associated proteins measured by the NP-MS workflow, using the model protein ∼ glucose + sex + age. H) Pearson linear modeling to identify fat percent associated proteins measured by the NP-MS workflow, using the model protein ∼ fat percent + sex + age.

### Protein associations with clinically relevant phenotypes

Previous studies report that glucose level and fat mass percentage change with advancing age and differ by sex in clinical studies [32-34].To identify associations between the circulating proteome and these phenotypes in mice, we performed a cross-sectional analysis of the clinical parameters of the 30-sample cohort. The demographics and phenotypic data are provided in **Supplementary Table 3.** Glucose and body composition are key age-associated clinical phenotypes and biomarkers of health outcomes and disease risk. In the cohort, glucose levels declined with age though did not show a significant relationship (**Fig. 5E**), while fat percentage was positively correlated with age (**Fig. 5F**). Linear modeling, adjusted for age and sex, revealed 43 proteins that were positively associated with glucose level, and 34 that were inversely associated (**Fig. 5G**). The top glucose associated proteins included Serpinf1 (**Supplementary Fig. 4A**), Myh10, and Cfhr1, among others. Similarly, linear modeling revealed that 162 proteins were positively associated with fat percentage, while 23 were inversely associated (**Fig. 5H**). The top fat percentage-associated proteins include Prcp (**Supplementary Fig. 4B**), Nt5e, Ubash3b, among others.

## DISCUSSION

Currently, there is a lack of tools available to comprehensively detect and quantify proteins in the low volumes of blood available from mouse studies, which generally cannot accommodate existing approaches such as extensive fractionation [35]. As a result, biomarker studies in mouse cohorts are sparse and unfeasible to most researchers. Furthermore, even with extensive processing, MS-based proteomic studies notably miss detection of a large portion of cytokines, a highly sought after subset of protein biomarkers with biological relevance across multiple domains of health and disease research. To our knowledge, the proteomics workflow reported in this study has enabled the most comprehensive survey of the serum proteome to date in low volume serum, enabling biomarker research in longitudinal mouse studies that include key biologically relevant proteins such as cytokines. The approach requires a total of 21 µL of serum: 20µL for the NP-processing MS workflow and 1 µL for the targeted cytokine assay.

A major technical challenge in blood proteomic studies of blood are the wide dynamic range of protein concentrations, spanning 10 to 12 orders of magnitude [13]. To address this, we employed a previously described automated and unbiased NP-based MS workflow [15, 17, 18, 29]. Leveraging physicochemically distinct nanoparticles, this workflow forms a ‘protein-corona’ at the blood-nanoparticle interface, effectively compressing the sample’s dynamic range. This uncovers low abundance proteins typically missed in standard blood analyses, enabling their detection by MS. The deep, quantitative, and consistent composition of the protein corona enables the use of this approach for human biomarker studies [15-18, 29, 36, 37]. Our study combined a five NP workflow with data independent acquisition (DIA) MS, to test the feasibility of adapting this approach to low-volume mouse studies, identifying almost 3992 proteins and 28,976 peptides in a study (n=30) using 20 µL samples, a 6 -fold improvement over the standard neat workflow. We also optimized a reproducible and quantitative pipeline for the analysis of protein-level changes from nanoparticle-fractionated peptide data. Although the greatest proteomic depth is achieved by combining 5 NP fractions, if fewer fractions are desired, the majority of the improvement in proteomic depth (>5-fold increase over neat samples) can be achieved by using only nanoparticle 2 (**Supplementary Fig. 2D**). Recently, two-NP kits have also shown comparable proteomic depth to five NP [7, 38, 39].

The NP-based workflow greatly enhanced the detection of biologically relevant proteins and candidate biomarkers. Cellular senescence and inflammation are key hallmarks of aging that have previously been associated with aging and various age-associated clinical phenotypes in humans [31, 40]. Compared with a standard workflow, which detects < 50 senescence-associated markers described by SenSig [26], the NP workflow identified > 200 SenSig proteins. The NP workflow also enabled the detection of low-abundance inflammatory proteins, such as CXCL12 and IL16. Despite these improvements, the MS-based workflow notably missed many widely studied cytokines often measured in biomarker studies, such as the IL6, a widely used clinical indicator of inflammation and a prototypical senescence-associated secreted protein [4]. Cytokines often go undetected in MS studies because they are at the low end of the dynamic range of blood and are generally small proteins with a small number of unique tryptic peptides that ionize well [20, 21]. Our data suggest that applying a mouse cytokine panel based on proximity extension assay technology [41] is highly complementary to the MS-based protocol, quantifying 39 cytokines undetected by MS, including IL6 in only 1 µL of additional serum. Standard antibody-based cytokine arrays would presumably be complimentary as well, although the sensitivity and volume requirements of other traditional technologies would differ. Overall, a workflow that combines both approaches, untargeted MS analysis and a targeted cytokine assay, provide a complementary and comprehensive set of biologically relevant biomarker candidates that are not accessible in low volumes by existing methods.

Enabled by this workflow, we identified unique and fundamental insights into age- and sex-dependent serum proteome in a subset of samples from the Study of Longitudinal Aging in Mice (SLAM), a deep and comprehensive longitudinal study of normative aging in mice that includes deep phenotyping and multi-omics [11]. Importantly, because the volume of blood collected in a study of this size and scope is limited, and must be portioned for multiple assays, it was critical that comprehensive proteomic measurements, and statistically significant associations with biological phenotypes, were possible within a 25 µL volume limit. Indeed, we observed significant insights into biological determinants of serum proteome composition, including age and sex. We found that Myosin light chain 3 (Myl3), a core muscle component known to increase in the serum during aging and muscle wasting [42, 43], was among the strongest positively age-associated proteins. Interestingly, one of the most negatively associated aging proteins, Paraoxonase 1 (Pon1), is an atheroprotective protein with a known age-associated decrease, and variants in this gene have been associated with longevity [44]. Importantly, the inclusion of three ages in this study highlights that most circulating proteins do not follow a linear trajectory throughout the lifespan, and completely different sets of proteins change into old age (24 months) versus geriatric (30 months). The changes we report in old mice highlight the degree of the enrichment of ‘inflammaging’ related changes (immune response, regulation of adaptive immunity, gliogenesis, etc.) that become apparent in old age, consistent with previous findings. However, a completely different set of functions including energetic and metabolic processes are changing in very old (geriatric) age, such as purine-containing compound metabolism, cellular nitrogen metabolism, ribose phosphate metabolism, precursor metabolites and energy, and NADH metabolism, among others. These findings demonstrate the sensitivity and robustness of this workflow to detect significant and even subtle changes in aging biology present at different life stages and suggest that future studies would benefit from the inclusion of multiple timepoints and longitudinal study designs. This automated workflow is scalable and is compatible with deeper longitudinal investigations including more mice and timepoints.

Few comparable published studies exist to draw comparisons with and validate the findings of this study. One notable study comprehensively surveyed the mouse proteome with extensive fractionation (3972 proteins in 128 fractions), identified numerous sex-associated proteins in the context of Alzheimer disease (AD) mouse model [35]. A second study performed targeted MS analysis of 375 plasma proteins in male and female mice from 30 knockout strains from the International Mouse Phenotyping Consortium, reported baseline sex differences with which to compare our data [45]. Remarkably, of all 33 sex-associated proteins reported both in our study and one or both existing studies, all exhibited the same directionality of sex-associations (**Supplementary Table 4**) [35, 45]. Several of these proteins, including complement components C8g, C6, C7, C8b, and C8a, lipocalins such as major urinary proteins-Mup20 and Mup3, serine protease inhibitors like Serpina1e and Serpina3k, Cysteine-rich secretory protein 1 (Crisp1), and the transmembrane glycoprotein Epidermal growth factor receptor (Egfr) were elevated in male mice in other studies [46-52]. In females, higher levels of proteins such as glycosylation-dependent cell adhesion molecule 1 (Glycam1), Adiponectin (Adipoq), cholinesterase (Bche), and Serpina6 were reported in other studies [53-56]. Importantly, pathway analysis revealed distinct sex-specific pathways in circulation, with strong female-specific changes in peptidase and glycosaminoglycan-related pathways (**Fig. 5d**), and a distinct set of membrane protein pathways in males (**Fig. 5c**). Altogether, these findings underscore the sexual dimorphism in blood proteins and the need to examine both sexes in the circulating proteome. Moreover, the consistent agreement of our results with reported protein changes in the literature further underscore the accuracy and robustness of the nanoparticle-MS workflow.

Blood glucose and fat percentage are widely collected clinical phenotypes that are associated with a broad range of health metrics and aging. We previously reported that higher glucose levels late in life are predictive of decreased mortality risk in SLAM [32]. To identify protein signatures that might be similarly associated with over health, metabolic state, and longevity, we evaluated serum proteins associated with glucose and body fat percentage. These indices can serve as physiological biomarkers to explore the intricate relationship between glucose homeostasis and body weight and adiposity to age [57]. Notably, we identified 77 and 185 proteins significantly associated with glucose level and fat percentage, respectively. The strongest positive correlate of glucose was Serpin family F member 1 (Serpinf1), a key protein in regulating glucose and insulin tolerance [58, 59]. Conversely, one of the strongest negative correlates of glucose, TBC1 Domain Family Member 17 (Tbc1d187is a key regulator of glucose uptake [60]. These results reveal that serum proteins are closely associated with two key age-related metabolic indices-glucose level and fat percentage- and may hold promise as biomarkers for predicting metabolic health and biological aging.

Limitations of this study include the small sample size, which may limit the generalizability of the biological findings, and the cross-sectional design that prevents the assessment of causal relationships over time. Future studies will address these limitations by incorporating larger sample sizes to enhance statistical power and generalizability, as well as adopting a longitudinal design to better capture temporal relationships and causal effects.

In summary, this study highlights the complementary strengths of two proteomic workflows: NP-based enrichment combined with MS, and the Proximity Extension Assay. Together, these approaches achieve comprehensive and quantitative profiling of the mouse serum proteome in low volumes, enabling blood biomarker studies in longitudinal mouse studies. The NP-based MS approach substantially broadened the proteome coverage, while PEA offered crucial sensitivity for detecting inflammatory cytokines. The minimal overlap between the two methods underscores their synergistic value, collectively enhancing the depth of circulating proteome analysis. This integrated strategy enabled to quantification of over 4,000 proteins, including key cytokines from mouse serum, providing novel insights into age-related changes, sex-specific differences, clinically relevant phenotypes. These findings contribute to a deeper understanding of the biology of aging and its translational relevance to human health. Looking ahead, we are optimistic that, when scaled, this quantitative workflow could significantly advance circulating proteome profiling, opening a new avenue for biomarker discovery and improved disease monitoring.

## METHODS

### Study Design

This study comprised two main phases, 1) optimization and benchmarking of sample preparation and LC-MS workflow in a feasibility study, and 2) biomarker discovery in an aging mouse cohort using an optimized proteomics workflow coupled with a bioinformatic analysis (**Fig.1**). For phase 1, the feasibility of detecting proteins in low volumes with the Proteograph workflow was assessed from a single pooled mouse sample. The quantitative benchmarking study was carried out similarly to previously described LFQ-Bench studies [28], using defined mixtures of plasma from bovine and human purchased commercially (Innovative Research) at five different ratios (n = 5 each). For phase 2, we assessed serum samples from 30 mice selected from SLAM at three different ages: young (12 months), old (24 months), and geriatric (30 months), with 10 mice chosen at each age group (50% female in each group). SLAM is an ongoing longitudinal cohort-based investigation of normative aging conducted at the National Institute on Aging (NIA) [11]. Mice were born at The Jackson Laboratory (Bar Harbor, ME), shipped to the NIA animal facility (Baltimore, MD) under a special breeding contract in cohorts of ∼200 12-week-old mice, and followed longitudinally thereafter. Animal husbandry, environmental factors, and diets are described elsewhere [61]. Briefly, Animal rooms were maintained at 22.2 ± 1°C and 30%–70% humidity. Mice were monitored twice daily for health status, and veterinary care was administered as necessary. Routine tests are performed to ensure that mice are pathogen-free and sentinel cages are maintained and tested according to the American Association for Accreditation of Laboratory Animal Care (AAALAC) criteria. All experimental protocols were conducted according to the guidelines and approved by the Animal Care and Use Committee of the National Institute on Aging, NIH, under protocol number 458-TGB-2027.

### Serum Preparation

Serum preparation from blood was performed in serum separator tubes (Sarstedt AG + Co. 41.1378.005). The serum was aliquoted and kept frozen at −80°C until proteomic analysis. Pooled mice serum was used for a feasibility study of workflow optimization.

### Glucose and Fat Mass Determination

Glucose levels in blood were measured from the submandibular vein in 6h fasted mice with the Contour Next EZ glucometer (Bayer). The body composition of unanesthetized mice, measuring percentage of fat mass was measured by nuclear magnetic resonance (NMR) using the Minispec LF90 (Bruker Optics) and normalized to body weight [62].

### Sample Preparation for Proteomic Analysis

All serum samples were processed with Seer Proteograph workflow using an SP100 automation instrument [15, 29]. Briefly, serum samples were diluted with TE buffer (10mM Tris, 1mM disodium EDTA, 150mM KCl) and 0.05% CHAPS and mixed with NP suspension, here each sample was incubated at 37 °C for 1 h with shaking at 300 rpm with five proprietary, physicochemically distinct nanoparticles for protein corona formation. Incubation allows high-affinity proteins to displace high-abundance proteins, resulting in a reproducible protein corona on each NP surface that probes the depth of the serum proteome. After incubation and magnetic separation, bound proteins were reduced, alkylated, and digested with Trypsin/ Lys-C using a digestion kit [15]. Peptides were purified by cleanup cartridge, and their concentration was measured by a quantitative colorimetric peptide assay kit from ThermoFisher Scientific (Waltham, MA), and finally lyophilized by vacuum dried and stored at -80 C until analyzed in a mass spectrometry.

### Data-independent Acquisition (DIA) - LC-MS/MS Analysis

Peptide samples were reconstituted with 10 μL loading buffer composed of 2% acetonitrile (ACN) and 0.1% formic acid (FA) in LC-MS grade water. Indexed retention time standards (iRTs, Biognosys) were spiked according to manufacturer’s instructions prior to MS analysis. All samples were analyzed using a Q Exactive HF-mass spectrometer (ThermoFisher Scientific) interfaced with an Ultimate 3000 RSLC nanoLC (ThermoFisher Scientific). Reverse phase LC was conducted with two mobile phases (A and B). Mobile phase A consisted of 0.1% FA, in water, and mobile phase B contained 0.1% FA, in ACN. Peptides were loaded onto nanoViper Acclaim PepMap 100 trap column (3 μm C_18_ particle, 75 μm × 2 cm ThermoFisher Scientific).

Then separated on a PepMap RSLC column (2 μm C_18_ particle, 75 μm × 500 mm,ThermoFisher Scientific). The trap and separation column were maintained at 45°C within the column oven and EASY-spray ionization source, respectively. The nanosource capillary temperature was set to 320 °C and the spray voltage was set to 2.0-2.4KV. For the feasibility and aging studies, peptides were separated using a 120 min linear gradient from 5-30% of solvent B at a constant flow rate 250 nL/min, with a total run time of 180 min. For quantitative benchmarking samples, a shorter 65 min gradient (2-30% of solvent B) was applied using same column and flow rate, with a total run time of 100 min.

All MS data were acquired in data-independent acquisition (DIA) mode. MS2 scans were configured to acquire 50 × 24 m/z DIA spectra (24 m/z isolation window) spanning the mass range 400-1000 m/z, staggering the DIA window placement by about half an isolation window (12.7 m/z) during alternating cycles as previously described [63]. MS1 scan spanning 400-1000 m/z was acquired at 60,000 resolution with an AGC target of 3-5e^6^ and a maximum injection time of 60-100 ms. MS2 DIA scans were acquired at 30,000 resolution with an AGC target of 1e^6^ and maximum injection time 50-70ms spanning the same range using 24 m/z windows with normalized collision energy of 27-30% used for precursor fragmentation in HCD mode. The acquisition of samples was block-randomized to avoid bias [64].

### MS Data Analysis

All raw MS files from the feasibility and aging studies were searched using the DIA-NN v1.8.1 algorithm within Proteograph Analysis Suite (PAS) v2.0. Peptide identification and quantification were performed with library-free mode searching MS/MS spectra against an in-silico generated spectral library of UniProtKB mouse FASTA database (UP000000589_10090). For quantitative benchmarking study, MS data were analyzed using the same pipeline with a combined database containing both human and bovine proteomes. The search parameters were set as follows: trypsin protease, 1 missed cleavage, fixed modification of Cys Carbamidomethylation, no Met oxidation, N-terminal Met excision, peptide length of 7-30 amino acids, precursor range of 300-1800 m/z, and fragment ion range of 200-1800 m/z. Peptides and Proteins FDR were set at 1% (Q-value). MS1 and MS2 mass accuracy was set to 10 ppm with Heuristic protein inference enabled. Peptide quantification was performed using a max representation (MaxRep) approach, where the single quantification value for a particular peptide represents the quantitation value of the nanoparticle most frequently measured across all samples with the highest intensity, and standard peptide-to-protein rollup was used on selected peptide values. During workflow optimization, for comparison, MaxLFQ [30] was also utilized for nanoparticle and peptide to protein rollup using a MaxLFQ R-script.

### Proximity Extension Assay

We used the Olink Target 48 Mouse Cytokine Proximity Extension assay (PEA) to explore a panel of 43 cytokines (Olink Proteomics) in the mouse serum samples. Assays were performed by AssayGate Inc. (Ijamsville, MD) according to manufacturer’s instructions.

### Statistical and Bioinformatic Analysis

In this study, serum samples were collected from a cohort that represents both sexes and multiple stages across the murine lifespan. A sufficient number of samples were included to detect statistically significant differences in key aging traits, as previously described in detail [62]. This allowed for identification of circulating protein associations with broad aging phenotypes. Raw precursor intensities were normalized to global median intensities. For Spearman and Pearson modeling, intensities were centered and scaled.

Bioinformatic analysis was conducted in RStudio Version 4.2.2. Expression levels of genes identified on the peptide and protein levels were compared between nanoparticle and rollup method using spearman correlation (stats package version 4.2.2). Pearson modeling was used to determine proteins and peptides associated with age and clinical traits within the cohort (stats package version 4.2.2). Pairwise expressional comparisons were made between young (12 months), old (24 months), and geriatric (30 months) mice using t-tests.

### Pathway enrichment and visualization

Overrepresentation and Gene Set Enrichment Analysis were conducted using the Cluster Profiler Package, version 4.6.0 [65].

### Data availability

All raw MS data files and associated quantitative data, metadata, and sequence databases used in this study are available on the MassIVE repository (dataset identifier: MSV000098399). FTP download link: ftp://massive-ftp.ucsd.edu/v10/MSV000098399/

## Acknowledgements

We gratefully acknowledge Lauren Brick and NIDA/NIA Visual Media for assistance in figure preparation. This work was supported by the National Institute on Aging (NIA) Intramural Research Program (IRP), NIH. N.B. was supported by a Geroscience Research Opportunities grant from Hevolution Foundation (HF-GRO-23-1199068-44. PI: Basisty). N.B. and R.d.C. were supported by a SenNet NIH Common Fund Grant (NIA U54 AG079779, PI: Elisseeff).

## Author Contribution

A.D., and M.B. performed the experiments. B.O. performed computational analysis and generated figures. A.D. was involved in MS data acquisition and analysis. A.D., B.O. and N.B. prepared the manuscript. M.F. helped with data management and editing. SLAM investigators, SC, NP, RC, N.B. provided the facilities and guidance for the study.

