## Supplementary Figures 1-4 for "Deep and Quantitative Proteomic Profiling of Low Volume Mouse Serum Across the Lifespan"

### Supplementary Figure 1: Feasibility Study

##### A) Protein groups and peptides identified across different sample volumes

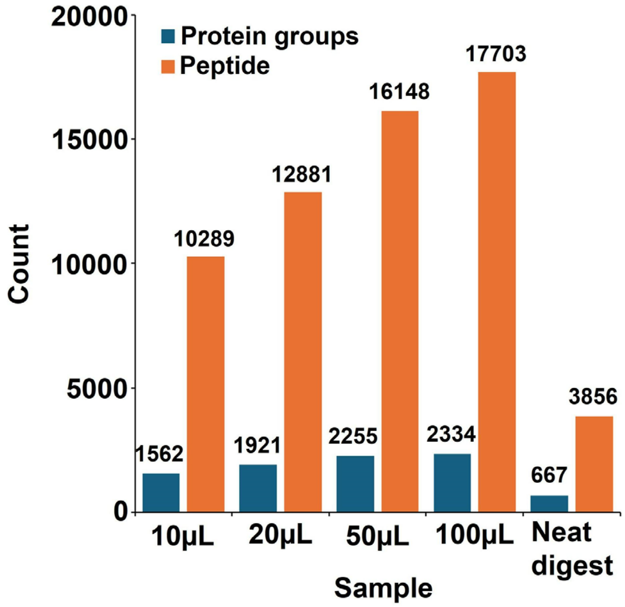


##### B) Protein groups identified using two serum preparation methods

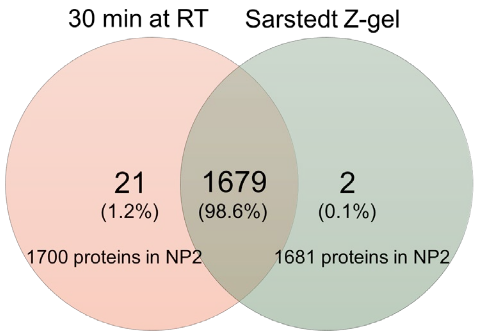


##### C) Candidate SASP proteins identified from 20 µL of serum

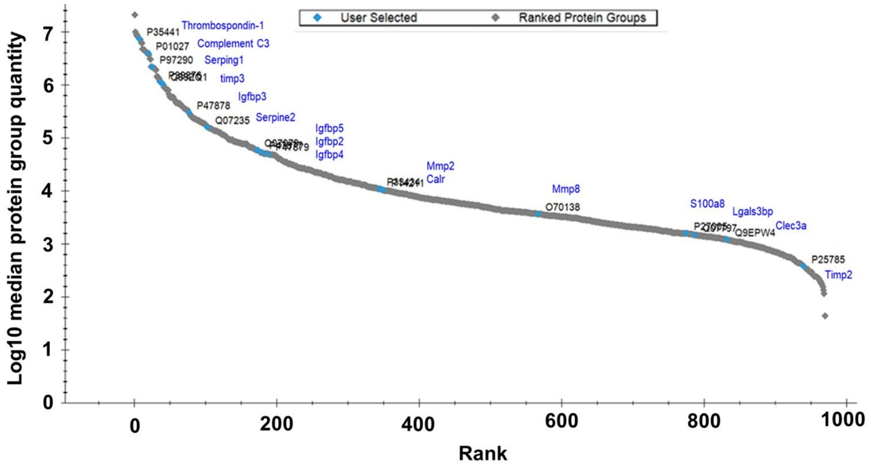


**Figure S1: A)** Number of protein and peptide groups identified in the feasibility study. **B)** Venn-diagram showing overlap of protein groups identified using two different serum preparation methods. **C)** Protein abundance ranked plot showing the dynamic range of protein detection

#### Supplementary Figure 2 : Comparison of Nanoparticle Rollup and Quantification

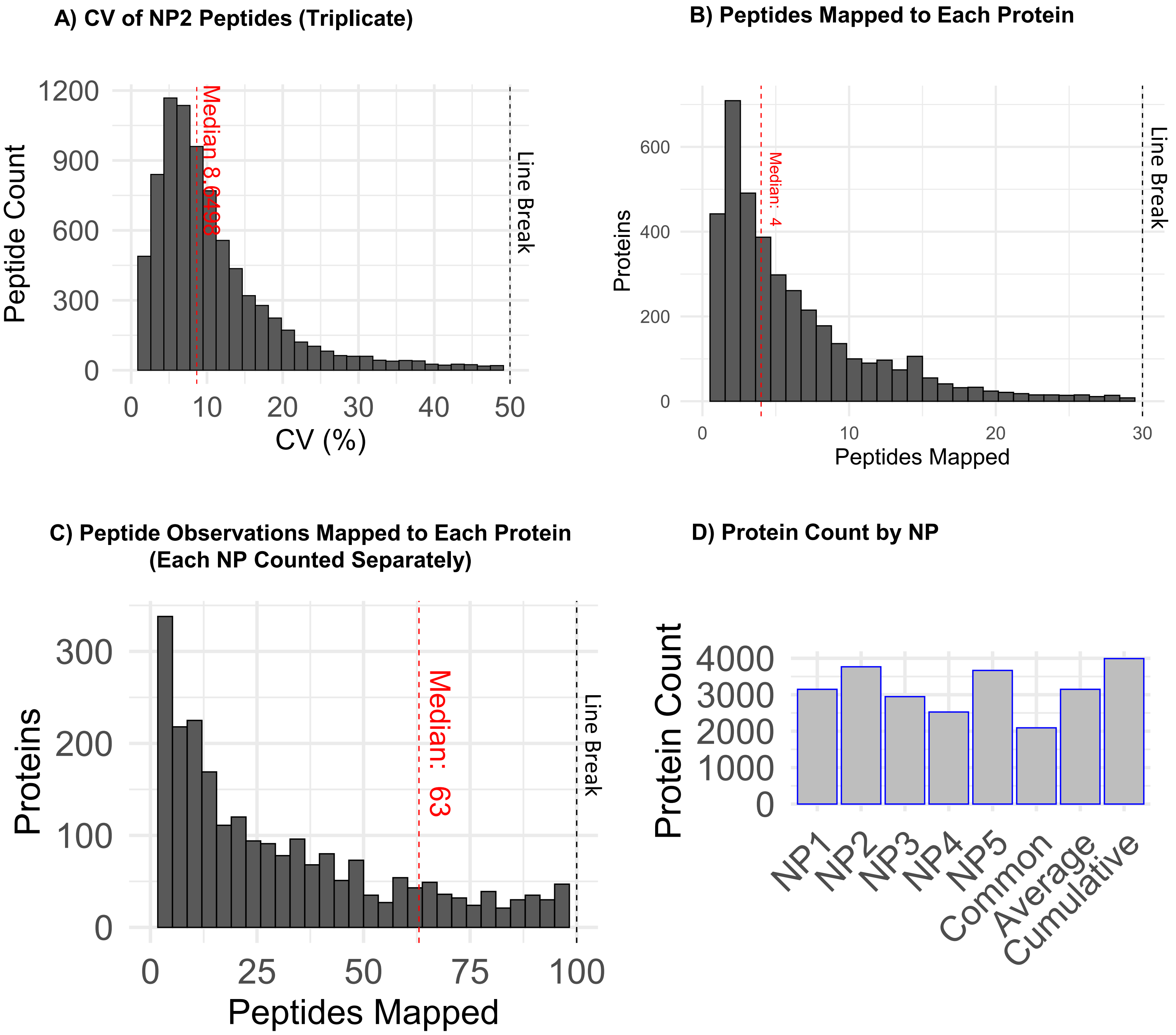


**Figure S2: A)** Coefficient of variation (CV=SD/Mean abundances * 100) of peptide level counts for NP2 across three technical replicates. Only peptides detected in all three samples were included.. For clarity, a line break at 50% is used. **B)** The number of peptides mapped to each protein in the pilot study. For clarity, a line break at 30 is used. **C)** The number of unique peptide observations mapped to each protein in the pilot study; with each NP counted separately. For clarity, a line break at 100 is used. **D)** The number of proteins identified by each NP in the pilot study.

#### Supplementary Figure 3: Sexual dimorphism

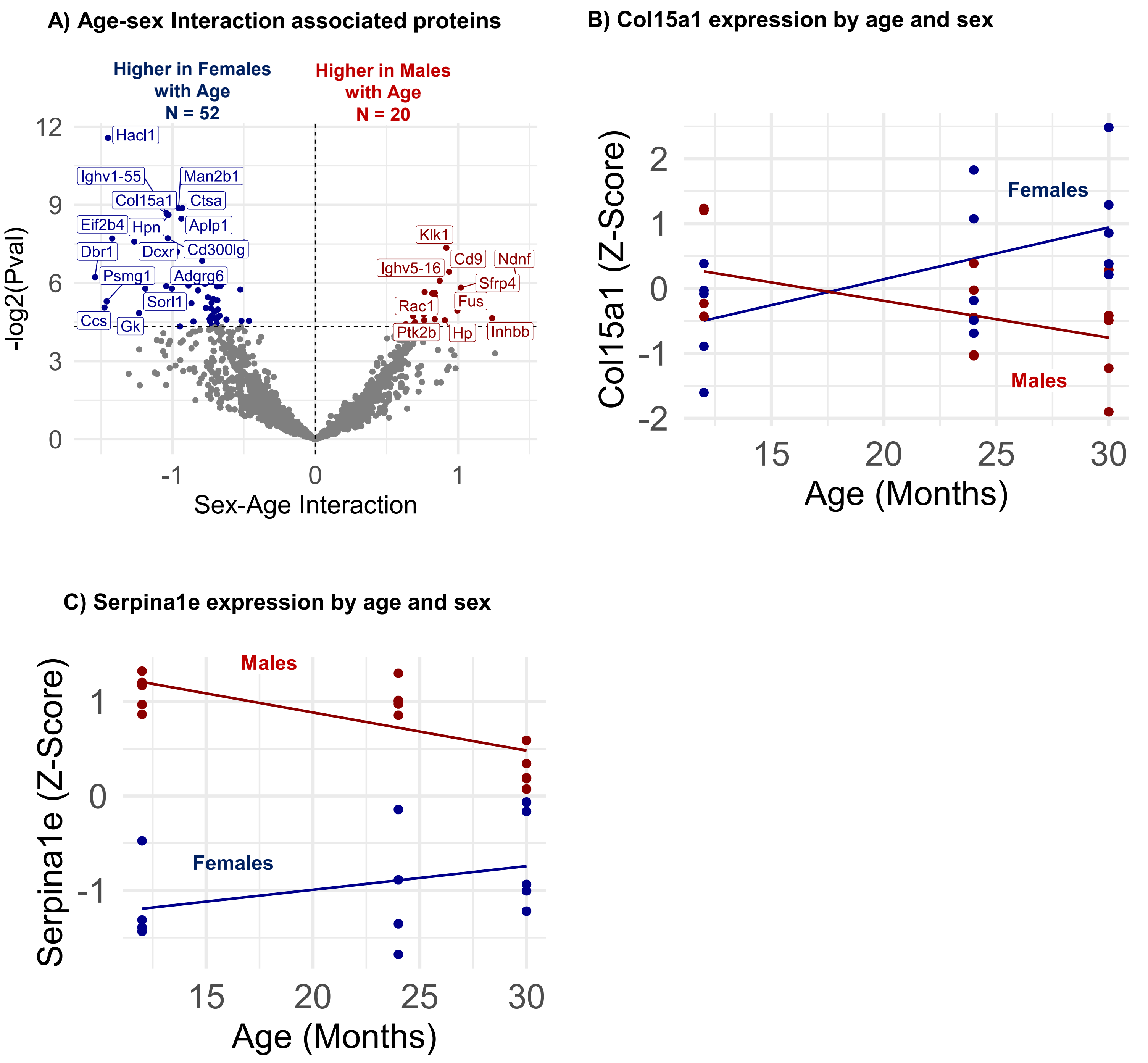


**Figure S3: A)** Pearson linear modeling to identify the interaction effects between sex and age on proteins measured by the NP-MS workflow, using the model: protein ~ sex * age. **B)** Col15a1 abundance by age and sex. **C)** Serpina1e abundance by age and sex.

**Supplementary Figure 4: Association of clinically relevant phenotypes**

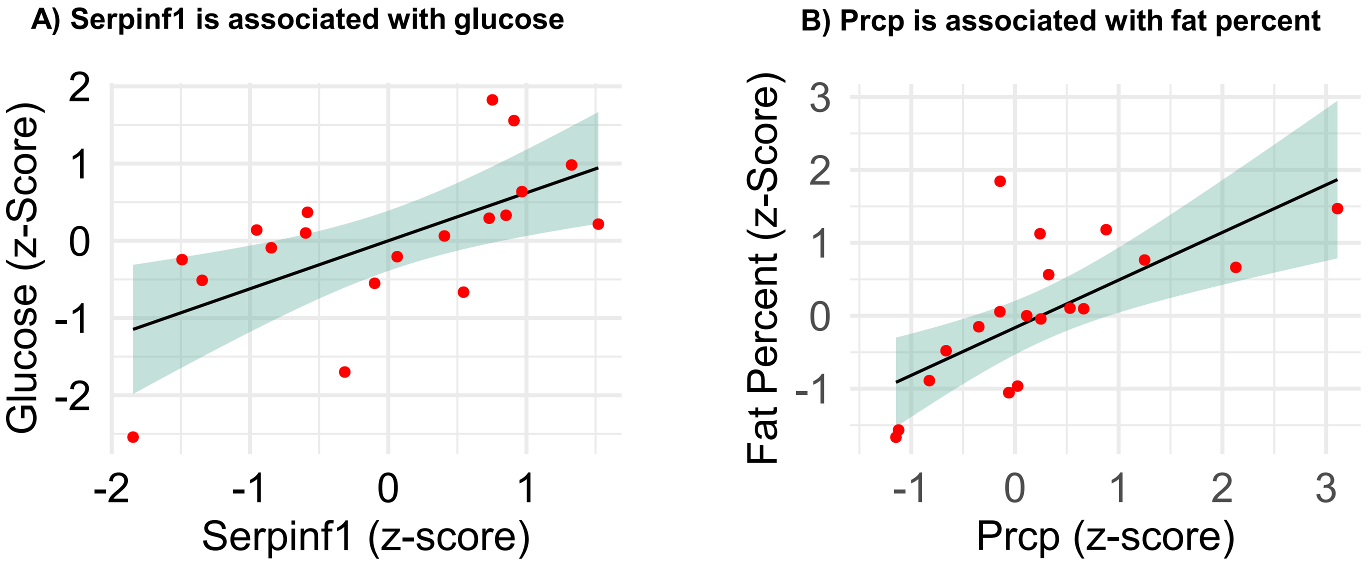


**Figure S4: A)** Serpinf1 abundance by glucose level in the pilot study cohort. **B)** Prcp abundance by fat percent in the pilot study cohort.
